# The landscape of structural variation in aye-ayes (*Daubentonia madagascariensis*)

**DOI:** 10.1101/2024.11.08.622672

**Authors:** Cyril J. Versoza, Jeffrey D. Jensen, Susanne P. Pfeifer

## Abstract

Aye-ayes (*Daubentonia madagascariensis*) are one of the 25 most critically endangered primate species in the world. Endemic to Madagascar, their small and highly fragmented populations make them particularly vulnerable to both genetic disease and anthropogenic environmental changes. Over the past decade, conservation genomic efforts have largely focused on inferring and monitoring population structure based on single nucleotide variants to identify and protect critical areas of genetic diversity. However, the recent release of a highly contiguous genome assembly allows, for the first time, for the study of structural genomic variation (deletions, duplications, insertions, and inversions) which are likely to impact a substantial proportion of the species’ genome. Based on whole-genome, short-read sequencing data from 14 individuals, >1,000 high-confidence autosomal structural variants were detected, affecting ∼240 kb of the aye-aye genome. The majority of these variants (>85%) were deletions shorter than 200 bp, consistent with the notion that longer structural mutations are often associated with strongly deleterious fitness effects. For example, two deletions longer than 850 bp located within disease-linked genes were predicted to impose substantial fitness deficits owing to a resulting frameshift and gene fusion, respectively; whereas several other major effect variants outside of coding regions are likely to impact gene regulatory landscapes. Taken together, this first glimpse into the landscape of structural variation in aye-ayes will enable future opportunities to advance our understanding of the traits impacting the fitness of this endangered species, as well as allow for enhanced evolutionary comparisons across the full primate clade.

## INTRODUCTION

Gaining a better understanding of the process of mutation is of fundamental importance to characterize genetic variation within and between populations and species, as well as to provide insights into both the drivers of local adaptation and the factors underlying disease. Over the past decades, the primary focus of many primate population genomic studies has been on elucidating the causes and consequences of point mutations, using single nucleotide variants (SNVs) to infer the rates and patterns of recombination, population demographic history, and natural selection (e.g., Auton et al. 2012; Simkin et al. 2014; Pfeifer and Jensen 2016; Stevison et al. 2016; Nielsen et al. 2017; Pfeifer 2017a, 2020a,b, 2021; Ghafoor et al. 2023; Johri et al. 2023; Soni et al. 2024a,b; Soni and Jensen 2024; Versoza et al. 2024a,b; Versoza, Lloret-Villas, et al. 2024). This common emphasis on SNVs, however, has resulted in a general neglect of the largest source of heritable variation, namely structural variation. Structural variants (SVs) – including copy number variants (defined here as deletions and duplications larger than 50 bp in size) and balanced rearrangements such as inversions – affect more nucleotides than SNVs in the primate genomes examined to date (Redon et al. 2006; Conrad et al. 2010; Pang et al. 2010; Sudmant et al. 2010, 2013, 2015a,b; Zarrei et al. 2015; Mao et al. 2024; and see the reviews of Conrad and Hurles 2007 and Gökçümen and Lee 2009). Moreover, due to their size, SVs often impact coding and regulatory regions which, in turn, can alter gene dosage, genome structure, or modify the timing and/or level of gene expression (Chaignat et al. 2011; Chiang et al. 2017), making SVs one of the main factors impacting phenotypic adaptation as well as disease susceptibility (Lin and Gökçümen 2019; and see reviews by Girirajan et al. 2011, Iskow et al. 2012, and Hollox et al. 2022).

Yet, despite their importance, the landscape of structural variation remains poorly characterized in many species. This neglect largely owes to the fact that SVs are more challenging to accurately identify and genotype than SNVs. On the one hand, the long read lengths of cutting-edge single-molecule sequencing technologies – in particular, Pacific Biosciences (15-20 kb at 99.95% accuracy; Olson et al. 2022) and Oxford Nanopore Technologies (10-100 kb at 99.26% accuracy according to the Q20+ Simplex Dataset Release) – facilitate the reliable discovery of SVs of different types and sizes; however, high costs and low throughput still prohibit the routine application of these technologies in many research areas. On the other hand, high-throughput short paired-end read sequencing (e.g., to 2 × 150 bp NovaSeq at 99.92% accuracy; Olson et al. 2022) tends to be more affordable but SV detection can be hampered by high false discovery rates, particularly in repetitive, complex, and highly polymorphic regions of the genome which are prone to errors in base calling and alignment from short-read data (Cameron et al. 2019; Kosugi et al. 2019; and see the discussion in Mahmoud et al. 2019).

Together with the progress in sequencing technology, a variety of short-read whole-genome callers have been developed that utilize different signals in the sequencing data to computationally detect SVs (see Supplementary Table S1 of Kosugi et al. 2019 for a summary of popular short-read SV callers). Typically, in assembly-free approaches, these signals include regional differences in read depth, changes in the direction and/or distance between read pairs (i.e., discordant read pairs), as well as unmatched read pairs that span SV breakpoints (i.e., split reads) (see Figure 2 in Alkan et al. 2011). Comprehensive benchmarking studies based on high-quality SV call sets obtained from deep-sequencing of human cell lines with multiple platforms (i.e., the *de facto* gold standard in the field) as well as simulated (ground truth) data have demonstrated that caller performance depends strongly on the SV type and size; as such, performance varies widely between approaches, with callers utilizing a combination of disparate read signals generally outperforming single-signal callers in terms of sensitivity (Cameron et al. 2017; 2019; Kosugi et al. 2019; Gabrielaite et al. 2021). For example, one of the largest benchmarking studies to date (Kosugi et al. 2019) showed that three of the best-performing short-read whole-genome SV callers – DELLY (Rausch et al. 2012), Lumpy (Layer et al. 2014), and Manta (Chen et al. 2016) – differ markedly in their precision and recall depending on the SV category.

Specifically, although Lumpy performed best for very small (50 - 100 bp) and small (100 bp - 1 kb) deletions (with mean precision / recall rates for whole-genome human resequencing data ranging from 76.9% / 13.1% to 90.3% / 26.2%) as well as inversions (40.2% / 0.7%), Manta exhibited a higher precision for medium-sized (1 - 100 kb) and large (100 kb - 1 Mb) deletions (93.3% / 27.0% and 36.9% / 8.3%) as well as small and medium-sized duplications (54.9% / 7.4% and 19.0% / 0.9%). In contrast, DELLY outperformed both Lumpy and Manta in the detection of large duplications (7.0% / 2.4%). Given the complementary strengths of different methodologies, the authors also explored multi-caller scenarios, demonstrating that precision for different SV types and sizes can be improved by applying a so-called "ensemble" approach that generates a call set based on SVs detected by several independent callers (see Supplementary Table S16 in Kosugi et al. 2019). This strategy is now widely employed in the field – though it should be noted that, unlike for SNVs (Pfeifer 2017b), no standardized best practices yet exist for SV discovery (see the discussion in Ho et al. 2020).

Methodologies aside, the great majority of work on the topic within primates to date has focused upon the great apes (Mao et al. 2024). However, in order to gain a broader evolutionary perspective, there would be great value in studying additional species from across the primate clade. One important evolutionary outgroup to the Haplorhini (which includes the apes), is the Daubentoniidae family of the Strepsirrhini suborder, which consists of the extinct giant aye-aye (*Daubentonia robusta*) (Nowak 1999) as well as extant aye-ayes (*Daubentonia madagascariensis*). Endemic to Madagascar, the world’s largest nocturnal primate (Kay and Kirk 2000) inhabits primary rainforests and dry undergrowth forest on the eastern, northern, and north-western parts of the island (Sterling 1994a; Louis et al. 2020) – however, widespread forest degradation, fragmentation, as well as slash-and-burn agriculture continues to destroy many of their native habitats (Suzzi-Simmons 2023), posing a severe threat to the survival of the species. Aye-ayes exhibit many distinct phenotypic traits (Sterling and McCreless 2006), including elongated, flexible middle fingers and rodent-like teeth that allow them to extract small insects from decaying wood (Erickson 1994). In addition to wood-boring insects, their diet includes a variety of seeds and fruits, making them important seed dispensers in their native forests (Sterling 1994b). Consequently, aye-ayes play a crucial role in maintaining the general health and balance of Madagascar’s flora and fauna. Yet, despite the aye-aye’s unique ecological and evolutionary significance, our knowledge of the population genetics of this elusive species remains limited (though see Perry et al. 2012, 2013 for insights into SNV diversity based on low-coverage sequencing data as well as Terbot et al. 2024). As one of the most critically endangered primate species on Earth (Louis et al. 2020), gaining insights into the structural variation landscape as a significant source of genetic diversity is thus vitally important for both conservation efforts of the species specifically, as well as to improve our understanding of the evolutionary history of primates in general.

Utilizing a high-precision ensemble approach for reliable SV discovery and genotyping from short-read sequencing data as described above, combined with previously developed methodology for SV curation in non-model organisms, we here analyze novel high-coverage genomic data of 14 individuals from ten trios (parents and their offspring) together with the recently released (Versoza and Pfeifer 2024), highly contiguous, well-annotated genome assembly for the species (a prerequisite for accurate SV discovery), to provide first insights into the genome-wide landscape of structural variation in this highly endangered primate.

## RESULTS AND DISCUSSION

SVs were detected by whole-genome sequencing of 14 aye-aye individuals from ten parent-offspring trios (Supplementary Figure 1) to an average autosomal coverage of 46.1x (range: 41.8x to 49.0x; Supplementary Table 1) – well above the 30x generally recommended for SV discovery and genotyping from short-read data (see Wold et al. 2021 and references therein). In brief, as SV detection can be hampered by high sequencing error rates and non-biological artefacts, raw reads were adapter and quality trimmed, before mapping them to the long-read genome assembly for the species and marking duplicates, which can result in spurious regions of extreme coverage (Supplementary Table 2). Based on these high-quality read mappings, SVs were then identified using an ensemble strategy that combined the strengths of local *de novo* assembly, with read depth, split-read, and discordant read approaches implemented in DELLY (Rausch et al. 2012), Lumpy (Layer et al. 2014), and Manta (Chen et al. 2016) – a methodology recently shown to result in robust and highly precise SV detection in humans (Subramanian et al. 2024). To increase precision, single-caller datasets were consolidated into a consensus call set of SVs identified by at least two of the three approaches using SURVIVOR (Jeffares et al. 2017) and subsequently filtered following the methodology described by Thomas et al. (2021) for another non-human primate species for which structural variation has been studied at the population-scale (rhesus macaque). In order to understand the potential medically-related impact of SVs, they were annotated using SnpEff (Cingolani et al. 2012), together with the gene annotations available from the aye-aye genome assembly (Versoza and Pfeifer 2024), and the putative relationship between large-effect SVs overlapping coding regions and diseases was assessed using the human database of Disease-Gene Associations with annotated Relationships among genes (eDGAR; Babbi et al. 2017) as a proxy.

A total of 1,133 autosomal SVs were identified in the 14 individuals, affecting 241,177 bp of the aye-aye genome (Figure 1). Of these 1,133 SVs, 1,000 were deletions (88.3%), 81 duplications (7.2%), 51 inversions (4.5%), and a single insertion – similar in proportion to the SV types previously observed in humans (89.4% deletions and 10.6% duplications; Brandler et al. 2016; *χ*2 = 0.009; df = 1, *p*-value = 0.9226) and rhesus macaques (88.3% deletions and 11.7% duplications; Thomas et al. 2021; *χ*2 = 0.016; df = 1, *p*-value = 0.8993). In concordance with these earlier studies (Brasó-Vives et al. 2020; Thomas et al. 2021), the majority of segregating deletions were short (median length: 172 bp; Figure 2a) – a pattern presumably resulting from the fact that deletions are often associated with deleterious fitness effects and are thus purged from the population, with purifying selection having been observed to be acting more strongly on longer deletions which more easily perturb protein function (Taylor et al. 2004; Itsara et al. 2010; Mills et al. 2011; Yang et al. 2024). Duplications and inversions tended to be longer (median duplication / inversion length: 424 bp / 1.1 kb; Figure 2a); thus, while being smaller in number, each event affected a larger proportion of base-pairs on average (Figure 2b). From a technical standpoint, this pattern reflects, at least in part, an ascertainment bias as duplications and insertions are more difficult to detect from short-read sequencing data than deletions (see discussions in Conrad and Hurles 2007; Sudmant et al. 2015b; Kosugi et al. 2019; Mahmoud et al. 2019; Delage et al. 2020).

**Figure 1.**
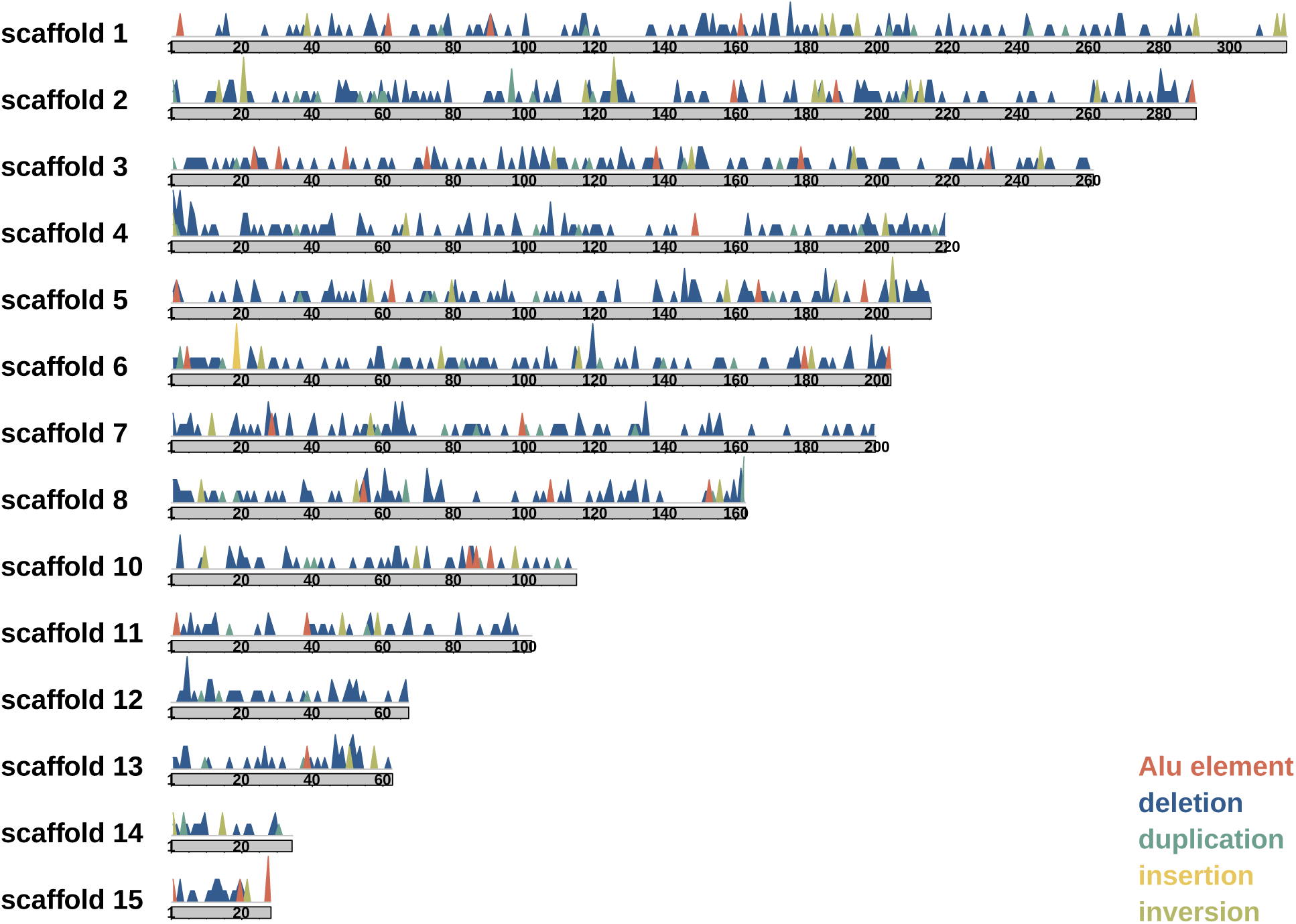
Landscape of structural variation in the aye-aye genome. Genome-wide map of structural variation (deletions are color-coded in blue, duplications in teal, insertions in yellow, and inversions in olive green) across autosomal scaffolds (note that scaffold 9, i.e., chromosome X, is not displayed), with peak height being proportional to the SV length. Putative *Alu* elements (shown in red) were removed prior to analyses.

**Figure 2.**
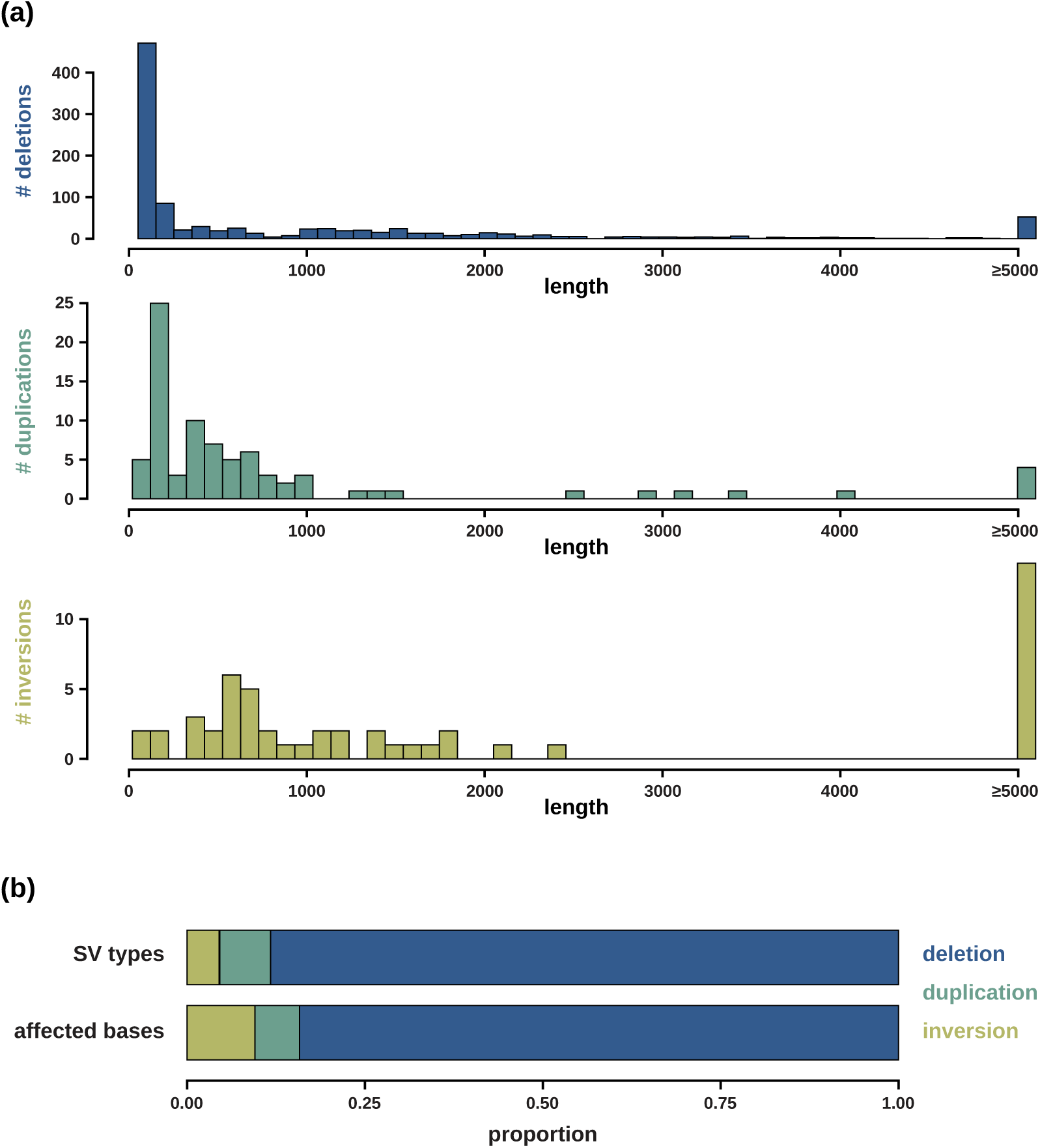
Characteristics of structural variation in the aye-aye genome. (a) Length distribution of structural variants (SVs; deletions are color-coded in blue, duplications in teal, and inversions in olive green; the single detected inversion is not shown). (b) Proportion of different SV types and base-pairs affected.

Per aye-aye individual, between 360 and 523 SVs were discovered (Supplementary Table 3) – a lower SV diversity than those previously observed in humans (Sudmant et al. 2015b) and rhesus macaques (Brasó-Vives et al. 2020; Thomas et al. 2021), consistent with the lower SNV diversity and effective population size of the species (Perry et al. 2012, 2013; Terbot et al. 2024). SVs were relatively evenly distributed across the genome (Supplementary Figure 2), with 12 SV-dense regions (≥ 10 SVs within a 10 Mb window) across the autosomal scaffolds (Supplementary Table 4). In concordance with observations in other primates (Bailey and Eichler 2006; Brasó-Vives et al. 2020; Thomas et al. 2021), these SV-dense regions were enriched in sub-telomeric parts of the genome which frequently harbor transposable elements that facilitate non-allelic homologous recombination – a biological process mediating structural variation (Conrad and Hurles 2007). Moreover, the number of SVs was strongly correlated with the length of the scaffold (deletion: *r* = 0.977, *p*-value = 2.24 x 10^-9^; duplications: *r* = 0.740, *p*-value = 0.0025; inversions: *r* = 0.798, *p*-value = 6.27 x 10^-4^; Supplementary Figure 3) as previously observed in other organisms (Thomas et al. 2021; Wold et al. 2022). In agreement with previous work in humans (Conrad et al. 2010; Belyeu et al. 2021), SVs were frequently harbored in gene-rich regions, with 36.8% and 20.4% residing within intronic and non-exonic, non-frameshift, non-missense genic regions, respectively. In addition to SVs within intergenic regions (41.6%), six SVs caused frameshifts, four SVs were located in exonic regions, three SVs were stop-related, and one SV resulted in a missense mutation (Supplementary Table 5). A total of 625 SVs (55.2%) were predicted to affect transcripts and a further 35 SVs (3.1%) were putative gene variants; the remainder were predicted to impact intergenic features. The vast majority of SVs were classified as modifiers (94.7%); the remaining SVs were predicted to have a high (4.0%), moderate (0.9%), and low (0.4%) impact (Figure 3), including several potential gene fusion events, exon losses, and frameshift mutations (Table 1). As expected, SVs with predicted high, moderate, and low effects were significantly enriched in genic regions (*χ*2 = 40.902; df = 1, *p*-value = 1.6 × 10^-10^). Out of the 45 major effect SVs, two deletions were located within disease-linked genes: (i) a frameshift variant in the cleavage factor polyribonucleotide kinase subunit 1 (CLP1) gene linked to pontocerebellar hypoplasia (PCH) subtype 10 – an autosomal recessive condition characterized by impaired brain development, motor neuron degeneration, and seizures (Karaca et al. 2014; Schaffer et al. 2014; and see review by van Dijk et al. 2018) – and (ii) a variant leading to a gene fusion in the opioid binding protein/cell adhesion molecule like (OPCML) gene – a tumor suppressor that is often epigenetically silenced in cancer, most prominently ovarian cancer (Birtley et al. 2019) (Table 1). Although the remaining major effect SVs were not predicted to exhibit a direct link to a disease, several ablated or disrupted genes, including those related to immune response (IGHV1-18 and IGHV1-24; Rodriguez et al. 2023) as well as circadian rhythm, particularly diurnal oscillations in light and temperature (BMAL2; Pando et al. 2001), were observed. Furthermore, several major effect SVs were located outside of coding regions and future work focusing on the potential regulatory impact of these changes would thus be of great interest.

**Figure 3.**
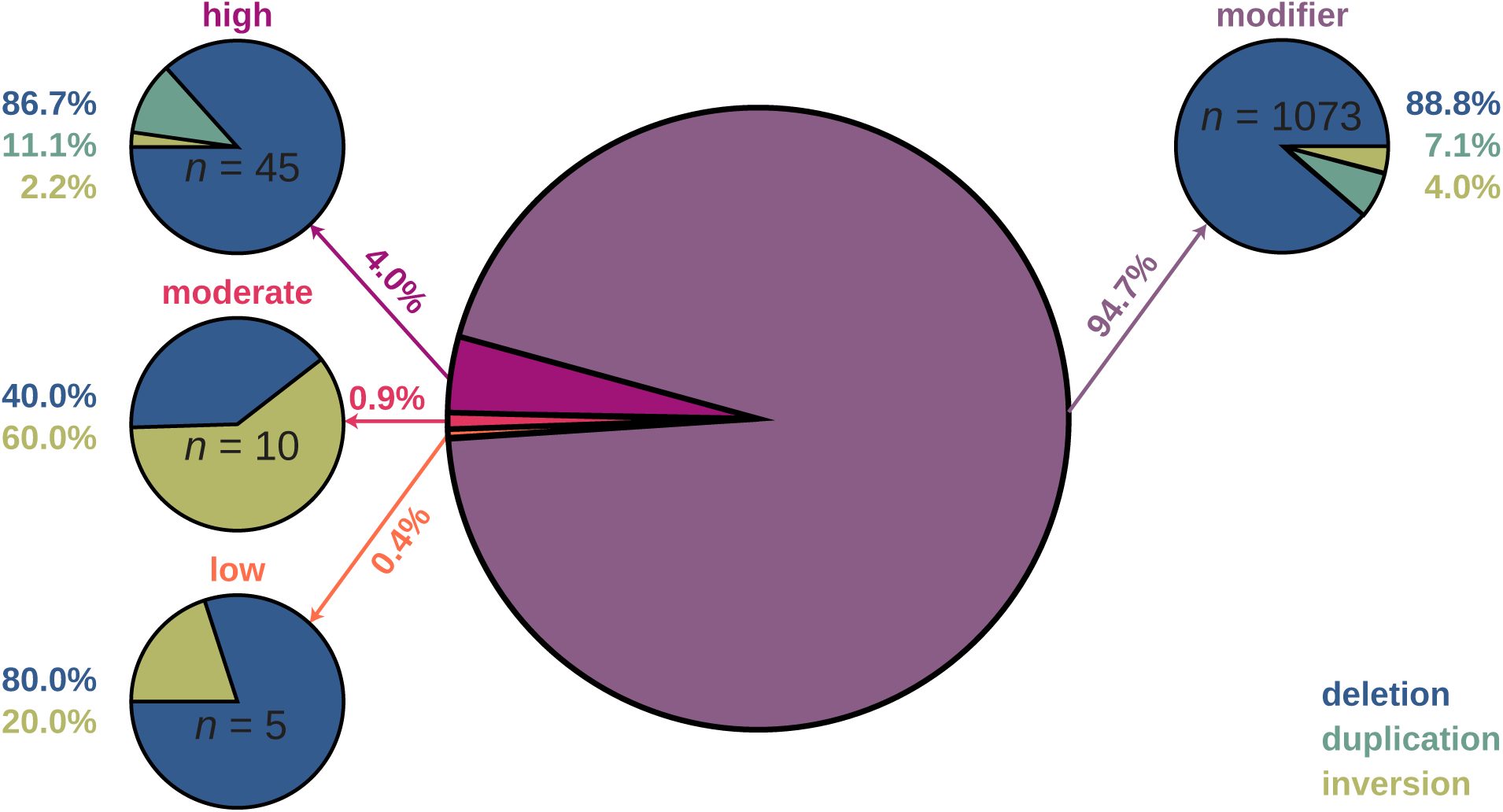
Annotation of structural variation in the aye-aye genome. The proportion of structural variants (deletions are color-coded in blue, duplications in teal, and inversions in olive green; the single detected inversion is not shown) classified as modifiers (shown in purple) as well as those predicted to have a high (pink), moderate (rose), and low (orange) impact.

**Table 1.** Structural variants with major effects in aye-ayes. . SVs in disease-linked genes are highlighted in red.

| scaffold | start | size | type | predicted effect | allele freq. | feature type | transcript type | eDGAR |
| --- | --- | --- | --- | --- | --- | --- | --- | --- |
| scaffold 1 | 114,706,693 | -1,678 | DEL | splice donor variant | 0.18 | transcript | pseudogene | N/A |
| scaffold 1 | 117,064,874 | 65,741 | DUP | stop gained | 0.04 | transcript | protein-coding | not a disease-linked gene |
| scaffold 1 | 117,231,258 | -3,148 | DEL | gene fusion | 0.46 | gene variant | – | not a disease-linked gene |
| scaffold 1 | 175,090,637 | -3,775 | DEL | gene fusion | 0.14 | gene variant | – | not directly linked to a disease |
| scaffold 1 | 206,727,921 | -888 | DEL | frameshift variant; splice acceptor & donor variant | 0.43 | transcript | protein-coding | disease-linked gene |
| scaffold 1 | 308,586,685 | -2,940 | DEL | gene fusion | 0.14 | gene variant | – | disease-linked gene |
| scaffold 2 | 193,565 | 65,573 | DUP | gene fusion | 0.36 | gene variant | – | not a disease-linked gene |
| scaffold 2 | 16,532,751 | -1,563 | DEL | splice acceptor variant | 0.32 | transcript | pseudogene | N/A |
| scaffold 2 | 59,474,585 | -1,613 | DEL | splice donor variant | 0.25 | transcript | pseudogene | N/A |
| scaffold 2 | 123,894,446 | -265 | DEL | splice donor variant | 0.43 | transcript | pseudogene | N/A |
| scaffold 2 | 147,696,112 | -1,992 | DEL | splice donor variant | 0.18 | transcript | pseudogene | N/A |
| scaffold 2 | 167,674,811 | -4,633 | DEL | transcript ablation | 0.36 | transcript | protein-coding | not directly linked to a disease |
| scaffold 2 | 280,664,663 | -352 | DEL | frameshift variant; splice acceptor & donor variant | 0.25 | transcript | protein-coding | not a disease-linked gene |
| scaffold 2 | 280,679,454 | -607 | DEL | splice acceptor & donor variant | 0.21 | transcript | protein-coding | not a disease-linked gene |
| scaffold 3 | 62,048,361 | -56 | DEL | frameshift variant | 0.32 | transcript | protein-coding | not a disease-linked gene |
| scaffold 3 | 114,478,661 | 24,999 | DUP | splice donor variant | 0.11 | transcript | pseudogene | not directly linked to a disease |
| scaffold 3 | 161,585,894 | -949 | DEL | splice acceptor variant | 0.54 | transcript | pseudogene | not directly linked to a disease |
| scaffold 3 | 195,072,212 | -1,137 | DEL | exon loss variant | 0.18 | transcript | pseudogene | not directly linked to a disease |
| scaffold 3 | 224,565,917 | -168 | DEL | splice acceptor variant | 0.18 | transcript | protein-coding | not directly linked to a disease |
| scaffold 3 | 226,807,718 | -93 | DEL | frameshift variant; splice donor variant | 0.04 | transcript | protein-coding | not a disease-linked gene |
| scaffold 3 | 240,213,435 | -54 | DEL | splice donor variant | 0.11 | transcript | protein-coding | not a disease-linked gene |
| scaffold 4 | 210,144,666 | -53,110 | DEL | splice acceptor & donor variant | 0.21 | transcript | protein-coding | not a disease-linked gene |
| scaffold 5 | 14,376,755 | -93 | DEL | splice donor variant | 0.18 | transcript | pseudogene | not directly linked to a disease |
| scaffold 5 | 18,775,900 | -3,663 | DEL | exon loss variant; splice acceptor & donor variant | 0.36 | transcript | protein-coding | not a disease-linked gene |
| scaffold 5 | 72,903,924 | 2,895 | DUP | stop gained | 0.39 | transcript | protein-coding | not directly linked to a disease |
| scaffold 5 | 80,139,609 | -167 | DEL | exon loss variant; stop lost; splice acceptor & donor variant | 0.18 | transcript | protein-coding | not directly linked to a disease |
| scaffold 5 | 94,879,702 | -34,807 | DEL | feature ablation | 0.54 | gene variant | – | not a disease-linked gene |
| scaffold 5 | 167,714,291 | -1,391 | DEL | gene fusion | 0.14 | gene variant | – | not a disease-linked gene |
| scaffold 5 | 170,514,034 | 480 | DUP | frameshift variant; splice acceptor & donor variant | 0.14 | transcript | protein-coding | not a disease-linked gene |
| scaffold 5 | 191,966,847 | -801 | DEL | splice donor variant | 0.07 | transcript | protein-coding | not a disease-linked gene |
| scaffold 6 | 146,740,803 | -66 | DEL | splice donor variant | 0.25 | transcript | pseudogene | not directly linked to a disease |
| scaffold 7 | 27,195,024 | -1,624 | DEL | gene fusion | 0.61 | gene variant | – | not a disease-linked gene |
| scaffold 7 | 40,345,354 | -1,612 | DEL | bidirectional gene fusion | 0.04 | gene variant | – | not directly linked to a disease |
| scaffold 7 | 164,612,242 | -210 | DEL | splice acceptor & donor variant | 0.21 | transcript | protein-coding | not a disease-linked gene |
| scaffold 7 | 188,327,797 | -58 | DEL | splice acceptor variant | 0.39 | transcript | pseudogene | not directly linked to a disease |
| scaffold 8 | 8,255,327 | 857 | INV | bidirectional gene fusion | 0.04 | gene variant | – | not directly linked to a disease |
| scaffold 8 | 55,672,556 | -1,536 | DEL | frameshift variant; splice donor variant | 0.21 | transcript | protein-coding | not a disease-linked gene |
| scaffold 8 | 55,677,369 | -163 | DEL | splice acceptor variant | 0.21 | transcript | protein-coding | not a disease-linked gene |
| scaffold 8 | 55,677,669 | -2,837 | DEL | exon loss variant; splice acceptor & donor variant | 0.21 | transcript | protein-coding | not a disease-linked gene |
| scaffold 10 | 84,664,633 | -90 | DEL | stop lost | 0.11 | transcript | protein-coding | not a disease-linked gene |
| scaffold 11 | 39,728,906 | -1,012 | DEL | splice acceptor & donor variant | 0.25 | transcript | protein-coding | not a disease-linked gene |
| scaffold 12 | 52,387,052 | -5,359 | DEL | splice acceptor & donor variant | 0.07 | transcript | pseudogene | not directly linked to a disease |
| scaffold 13 | 26,376,628 | -238 | DEL | splice acceptor & donor variant | 0.43 | transcript | protein-coding | not a disease-linked gene |
| scaffold 13 | 26,376,969 | -990 | DEL | splice acceptor & donor variant | 0.43 | transcript | protein-coding | not a disease-linked gene |
| scaffold 13 | 51,852,118 | -42,903 | DEL | feature ablation | 0.11 | gene variant | – | not a disease-linked gene |

The three-generation pedigree structure of this study also offered an opportunity to study Mendelian inheritance in the ten parent-offspring trios. Out of the 1,133 identified SVs, 114 sites (10.1%, including 99 deletions, eight duplications, and seven inversions) exhibited genotype-based Mendelian inconsistencies and were thus independently visualized for validation – a strategy previously shown to be in agreement with other orthogonal validation techniques such as ddPCR and long-read sequencing (Bertolotti et al. 2020; Belyeu et al. 2021). Perhaps unsurprisingly, given the orders of magnitude lower rates of structural mutation compared to point mutation previously observed in other primates (Werling et al. 2018), the lower genetic diversity of aye-ayes compared to the great apes (Perry et al. 2012, 2013; Terbot et al. 2024), and the small number of individuals in the cohort, no genuine *de novo* SVs were detected.

In addition to the small pedigree preventing the detection of rare SVs, a general caveat of short-read approaches, such as the ones employed here, is their inability to accurately identify SVs in low-complexity, highly repetitive regions of the genome (Chaisson et al. 2019). Due to the high false discovery rates frequently observed in such regions (Cameron et al. 2019; Kosugi et al. 2019; Mahmoud et al. 2019), regions harboring gaps, repeats, and/or extreme coverage were excluded from this study to increase precision. Additionally, stringent filtering was applied to the remaining regions to limit the dataset to SVs of high confidence and avoid spurious calls. It should be noted, however, that such an exclusion and filtering will necessarily lead to an underestimation of SVs, particularly those driven by homology-mediated mechanisms such as non-allelic homologous recombination, fork stalling and template switching, and microhomology-mediated break-induced replication (Belyeu et al. 2021).

In order to assess the SV calling and genotyping accuracy of the employed ensemble approach, the 1,596 genotypes at the 114 sites flagged as Mendelian violations were manually inspected in the pedigree. Interestingly, out of the curated sites, 35 (30.7%) were fixed for the reference allele (i.e., there was no read support for a SV at the site) and six (5.3%) were fixed for the alternative allele (Supplementary Table 6). Moreover, in contrast to a previous simulation study which observed high genotyping precision for deletions (91.9–94.1%), duplications (59.7–84.3%), inversions (87.8– 88.0%), and insertions (97.7%) in all three callers (Supplementary Table S15 in Kosugi et al. 2019), visual curation of 1,596 genotypes at the sites of Mendelian violations in the pedigreed dataset revealed a high rate of genotyping error (35.7%; Supplementary Table 6, and see Supplementary Figure 4 for an example of an incorrectly called genotype detected during the manual review). Taken together, these observations highlight that accurate SV calling and genotyping based on short-read data remains challenging (see also the discussion in Sibbesen et al. 2018), and emphasizes the importance of manual curation in studies of structural variation.

## CONCLUSION

With fewer than 1,000 to 10,000 individuals estimated to remain in the wild, aye-ayes are imminently threatened by extinction. Gaining insights into the genetic diversity of the species is thus of vital importance, and the first view of the landscape of structural variation presented here will be crucial to advance our understanding of the connection between genotypes and phenotypic traits relevant to conservation efforts and species recovery. In addition, as an important outgroup to the Haplorhini, this genomic data will allow for deeper comparative analyses across the primate clade to further our understanding of primate evolutionary history. In this regard, it is important to keep in mind that, although many similarities emerged between the structural variant landscape of aye-ayes and those of other primates studied to date, SV discovery approaches can have large impacts on the accuracy of both SV calls and their genotypes, thus hindering quantitative comparisons across studies. Moving forward, to facilitate meaningful comparisons, an important emphasis of future comparative genomic studies should thus be on the development of streamlined, uniform pipelines across the primate clade. Moreover, as short-read approaches are biased with regards to the SV types and sizes that they are able to detect, future studies should, whenever feasible, complement short-read data with long-read and/or optical sequencing approaches to obtain insights into the full spectrum of structural variation, including translocations and complex SVs. Ultimately, there is a pressing need to combine novel genomic resources, such as the one presented here, with ecological and evolutionary research in order to aid the development of more effective conservation strategies for this charismatic species.

## MATERIALS AND METHODS

### Animal subjects

This study was approved by the Duke Lemur Center’s Research Committee (protocol BS-3-22-6) and Duke University’s Institutional Animal Care and Use Committee (protocol A216-20-11). The study was performed in compliance with all regulations regarding the care and use of captive primates, including the U.S. National Research Council’s Guide for the Care and Use of Laboratory Animals and the U.S. Public Health Service’s Policy on Human Care and Use of Laboratory Animals.

### Sample collection, preparation, and sequencing

Genomic DNA was extracted from peripheral blood samples of 14 aye-aye (*D. madagascariensis*) individuals (six males and eight females) originating from a single three-generation pedigree using the PureLink Genomic DNA Mini Kit (ThermoFisher Scientific, Waltham, MA, USA) and quantified using a Qubit 2.0 Fluorometer (ThermoFisher Scientific, Waltham, MA, USA). Following manufacturer’s instructions, a sequencing library was prepared for each sample using the NEBNext^®^ Ultra^TM^ II DNA PCR-free Library Prep Kit (New England, Ipswich, MA, USA). Quality control of each library was performed using a High Sensitivity D1000 ScreenTape on an Agilent TapeStation (Agilent Technologies, Palo Alto, CA, USA). Libraries were quantified using a Qubit 2.0 Fluorometer (ThermoFisher Scientific, Waltham, MA, USA) and real-time PCR (Applied Biosystems, Carlsbad, CA, USA). Each library was paired-end sequenced (2 × 150 bp) on an lllumina NovaSeq platform (Illumina, San Diego, CA, USA).

### Read mapping

FastQC v.0.11.9 (https://www.bioinformatics.babraham.ac.uk/projects/fastqc/) and Cutadapt v.1.18 (https://cutadapt.readthedocs.io/en/stable/) embedded within TrimGalore v.0.6.10 (https://github.com/FelixKrueger/TrimGalore) were used to trim low-quality bases (with a Phred score < 20) and remove Illumina adapter sequences from the 3’-ends of the reads as they can lead to incorrect mappings. Afterward, the quality-controlled reads were mapped to the chromosome-level genome assembly for the species (DMad_hybrid; GenBank accession number: JBFSEQ000000000; Versoza and Pfeifer 2024) using BWA-MEM v.0.7.17 (Li and Durbin 2009). Read mappings were sorted, duplicates marked, and indexed using SAMtools *sort* v.1.20 (Danecek et al. 2021), GATK4 *MarkDuplicates* v.4.5 (Van der Auwera and O’Connor 2020), and SAMtools *index* v.1.20, respectively.

### Quality control

SV detection can be hampered by high sequencing error rates, uneven read coverage, and/or skewed insert size distributions, thus the quality of the read mappings and coverage distributions for each individual were assessed using SAMtools v.1.16 (Danecek et al. 2021) and goleft v.0.2.6 (https://github.com/brentp/goleft) prior to variant calling. Moreover, as regions harboring gaps, repeats, and/or extreme coverage frequently lead to mapping errors (Mahmoud et al. 2019), such genomic regions were excluded during the variant calling. In brief, sample coverage was estimated with mosdepth v.0.3.8 (Pedersen and Quinlan 2018) and high-coverage regions (defined here as regions exhibiting more than 10-fold of the mean autosomal coverage) as well as repetitive regions (including retroelements, DNA transposons, simple repeats, and low-complexity repeats) annotated in the aye-aye genome assembly (Versoza and Pfeifer 2024) were excluded.

### SV calling and genotyping

To increase precision, autosomal SVs were jointly called in the 14 aye-aye individuals using three of the best-performing short-read whole-genome SV callers according to recent benchmarking studies (Kosugi et al. 2019; Gabrielaite et al. 2021): DELLY v.1.2.6 (Rausch et al. 2012), Manta v.1.6.0 (Chen et al. 2016) and Lumpy v.0.2.13 (Layer et al. 2014) embedded within Smoove v.0.2.6 (https://github.com/brentp/smoove).

DELLY uses a combination of paired-end, read depth, and split-read signals for SV discovery. DELLY *call* was used to detect SVs from the read mappings, excluding (*-x*) repetitive and high-coverage regions as detailed above. Low-quality (*LowQual*) calls with fewer than three paired-end (*PE*) reads supporting a variant or with a mean mapping quality of less than 20 were discarded using BCFtools *view* v.1.10.2 with the *-e ’ FILTER=="LowQual" || FORMAT/FT=="LowQual" ’* flag, limiting the call set to precise SVs with split-read support at nucleotide resolution.

Manta combines paired- and split-read signals to detect, assemble, and genotype SVs. A configuration file was created (using the built-in *configManta.py* script) that provides information on the samples (*--bam*) and reference assembly (*--referenceFasta*) before running the two-step workflow (*runWorkflow.py*), consisting of a genome scan to identify candidate regions, followed by SV discovery, breakend assembly, genotyping, and filtering. Reported inversions were reformatted into single inverted sequence junctions using the built-in *convertInversions.py* script. SV calls were limited to variants outside of repetitive and high-coverage regions that passed all filter criteria using VCFtools *--exclude-bed* v.0.1.14 (Danecek et al. 2011) and BCFtools *view* v.1.10.2 with the *-i ’ FILTER=="PASS" ’* option, respectively. In brief, these filters excluded low-quality SVs (QUAL < 20 and, for those smaller than 1kb, sites where the proportion of reads in all individuals with a MAPQ0 around the breakend exceeds 40%), SVs larger than the paired-end fragment size without paired-end read support for the alternative allele, deletions and duplications inconsistent with diploid expectations, as well as SVs with breakends occurring in regions of excessive read depth (defined here as more than three times the median chromosome depth).

Lumpy utilizes regional differences in read depth to identify SVs; in addition, Lumpy detects unmatched read pairs by extracting split-read alignments (using the built-in *extractSplitReads_BwaMem* script) from discordant paired-end alignments (obtained using the ’ *samtools view -b -F 1294* ’ command). To accelerate the Lumpy workflow, the Smoove wrapper script *call* was used to parallelize these different steps, calling SVs outside of problematic regions (*--exclude*) and directly genotyping (*--genotype*) detected SVs using the Bayesian likelihood genotyper SVTyper v.0.7.0 (Chiang et al. 2015). By default, Smoove implements a series of filters that remove spurious alignments and improve specificity. Specifically, Smoove excluded reads that were soft-clipped at both ends, contained more than three mismatches, or exhibited alternative matches. To avoid spurious calls, Smoove further discarded split-reads for which the reads in a pair mapped to different chromosomes, split or discordant reads with a high depth of coverage (> 1000) as well as orphaned reads (i.e., reads without a mate). Following the developer’s recommendations (https://github.com/brentp/smoove), calls were annotated using *smoove annotate* and limited to sites with high-quality heterozygotes (i.e., SVs with a mean Smoove heterozygote quality [MSHQ] score larger than 3). Additionally, deletions and duplications were limited to sites with a fold-change of variant depth relative to flanking regions (DHFFC) of less than 0.7 and relative to genomic regions with similar GC-content (DHBFC) larger than 1.3, respectively.

In order to obtain high-precision calls, individual, single-caller datasets were consolidated into a consensus call set of SVs identified by at least two of the three approaches using SURVIVOR *merge* v.1.0.7 (Jeffares et al. 2017), merging any SVs of the same type that are closer than 500 bp.

### SV filtering

To reduce false positives, the consensus call set was filtered following the methodology described by Thomas et al. (2021). In brief, SVs present in all or all but one individual were removed as these are likely the result of local mis-assembly. Furthermore, SVs larger than 100 kb as well as those of low quality (QUAL < 100) were excluded to further limit the number of spurious variants in the dataset. The remaining SVs were then annotated with read depth information using Duphold v.0.2.1 (Pedersen and Quinlan 2019) embedded within Smoove v.0.2.6, and deletion and duplication events were limited to those exhibiting a fold-change of coverage of <0.7 and >1.3, respectively. Lastly, putative *Alu* mobile element insertions were filtered out by removing any SVs with a length between 275 and 325 bp.

### Functional annotation

SVs were annotated using SnpEff v.5.2 (Cingolani et al. 2012) based on gene annotations available in the aye-aye genome assembly (Versoza and Pfeifer 2024). In order to understand the potential medically-related impact of SVs, the putative relationship between large-effect SVs overlapping coding regions and diseases was assessed using the database of Disease-Gene Associations with annotated Relationships among genes (eDGAR; Babbi et al. 2017), with information from the Online Mendelian Inheritance in Man (OMIM; Amberger et al. 2017), humsavar (UniProt Consortium et al. 2015), and ClinVar (Landrum et al. 2016) databases embedded within.

### Identification of *de novo* SVs and assessment of SV calling / genotyping accuracy

Based on the final SV call set, Mendelian violations were identified using BCFtools v.1.20 (Danecek et al. 2021) with the +*mendelian* plugin and visually reviewed using Samplot v.1.3.0 (Belyeu et al. 2021) to identify *de novo* SVs and assess SV calling / genotyping accuracy.

## ACKNOWLEDGEMENTS

We would like to thank Erin Ehmke, Kay Welser, and the Duke Lemur Center for providing the aye-aye samples used in this study, and Fritz Sedlazeck as well as members of the Jensen Lab and Pfeifer Lab for helpful discussions. DNA extraction, library preparation, and Illumina sequencing was conducted at Azenta Life Sciences (South Plainfield, NJ, USA). Computations were performed on the Sol supercomputer at Arizona State University (Jennewein et al. 2023). This is Duke Lemur Center publication # XXXX.

