## Supplementary Materials for "The landscape of structural variation in aye-ayes (*Daubentonia madagascariensis*)"

| ID (pedigree) | sex | coverage |
|---|---|---|
| 1 | male | 45.3 |
| 2 | female | 43.9 |
| 3 | male | 48.8 |
| 4 | female | 46.0 |
| 5 | male | 49.0 |
| 6 | male | 48.6 |
| 7 | male | 48.7 |
| 8 | female | 48.1 |
| 9 | male | 46.4 |
| 10 | female | 46.6 |
| 11 | female | 45.9 |
| 12 | female | 41.8 |
| 13 | female | 44.8 |
| 14 | female | 44.2 |

**Supplementary Table 1**. Samples and autosomal coverage after quality-control.

| ID (pedigree) | % mapped reads | % properly paired reads |
|---|---|---|
| 1 | 99.9 | 98.6 |
| 2 | 99.8 | 99.1 |
| 3 | 99.9 | 98.9 |
| 4 | 99.9 | 99.2 |
| 5 | 99.9 | 98.6 |
| 6 | 99.9 | 98.6 |
| 7 | 99.9 | 98.1 |
| 8 | 99.9 | 98.5 |
| 9 | 99.9 | 98.3 |
| 10 | 99.9 | 98.8 |
| 11 | 99.9 | 98.6 |
| 12 | 99.9 | 99.4 |
| 13 | 99.9 | 99.2 |
| 14 | 99.9 | 98.9 |

**Supplementary Table 2**. Summary statistics of read mappings.

| ID (pedigree) | # SVs |
|---|---|
| 1 | 367 |
| 2 | 360 |
| 3 | 392 |
| 4 | 377 |
| 5 | 487 |
| 6 | 479 |
| 7 | 492 |
| 8 | 488 |
| 9 | 523 |
| 10 | 512 |
| 11 | 488 |
| 12 | 410 |
| 13 | 424 |
| 14 | 456 |

**Supplementary Table 3**. Number of structural variants (SVs) identified per individual.

| scaffold | region | # SVs |
|---|---|---|
| 1 | 150–160 Mb | 10 |
| 2 | 120–130 Mb | 11 |
| 2 | 280–290 Mb | 11 |
| 4 | 0–10 Mb | 20 |
| 4 | 190–200 Mb | 10 |
| 5 | 200–210 Mb | 11 |
| 6 | 0–10 Mb | 10 |
| 7 | 60–70 Mb | 11 |
| 8 | 0–10 Mb | 10 |
| 13 | 50–60 Mb | 10 |
| 14 | 0–10 Mb | 12 |
| 15 | 10–20 Mb | 12 |

**Supplementary Table 4**. Structural variant (SV) dense regions.

| # SVs | annotation |
|---|---|
| 471 | intergenic |
| 417 | intronic |
| 231 | non-exonic, non-frameshift, non-missense genic |
| 6 | frameshift |
| 4 | exonic |
| 3 | stop-related |
| 1 | missense |

**Supplementary Table 5**. Structural variant (SV) annotations.

| SV | | | | | individual | | | | | | | | | | | | | |
|---|---|---|---|---|---|---|---|---|---|---|---|---|---|---|---|---|---|---|
| **type** | **scaffold** | **start** | **size** | **freq** | **1** | **2** | **3** | **4** | **5** | **6** | **7** | **8** | **9** | **10** | **11** | **12** | **13** | **14** |
| **deletions** | scaffold 1 | 60,617,712 | 80 | 0.00 | 0/0 | 0/0 | 0/0 | 1/1 (0/0) | 0/0 | 0/0 | 0/0 | 0/0 | 0/0 | 1/1 (0/0) | 0/0 | 0/0 | 0/0 | 1/1 (0/0) |
| | scaffold 1 | 116,155,993 | 1,486 | 0.36 | 0/0 | 1/1 | 0/0 | 1/1 (0/1) | 0/1 | 0/1 | 0/0 | 0/1 | 0/1 | 1/1 (0/1) | 0/1 | 0/0 | 1/1 (0/1) | 0/0 |
| | scaffold 1 | 136,546,146 | 113 | 0.36 | 0/1 | 0/0 | 0/0 (0/1) | 0/0 | 0/1 | 0/1 | 0/0 | 0/1 | 0/0 (0/1) | 0/0 (0/1) | 0/0 | 0/1 | 0/0 | 1/1 |
| | scaffold 1 | 167,616,947 | 78 | 0.18 | 0/0 | 1/1 (0/1) | 0/0 | 1/1 (0/1) | 0/0 | 0/0 | 1/1 (0/1) | 0/0 | 0/0 | 0/1 | 0/0 | 0/0 | 1/1 (0/1) | 0/0 |
| | scaffold 1 | 179,207,030 | 2,260 | 0.21 | 0/0 | 0/0 | 0/1 | 0/0 | 0/0 | 0/0 | 1/1 (0/1) | 0/0 | 0/1 | 1/1 (0/1) | 0/0 | 1/1 (0/1) | 0/0 | 0/1 |
| | scaffold 1 | 180,389,032 | 1,428 | 0.36 | 1/1 | 1/1 | 0/0 | 0/0 | 1/1 | 1/1 | 0/0 | 1/1 (0/1) | 0/0 | 0/0 | 0/0 | 0/0 | 1/1 (0/1) | 0/0 |
| | scaffold 1 | 191,639,816 | 106 | 0.00 | 1/1 (0/0) | 0/0 | 0/1 (0/0) | 0/0 | 0/0 | 0/0 | 1/1 (0/0) | 0/0 | 0/0 | 1/1 (0/0) | 0/0 | 1/1 (0/0) | 1/1 (0/0) | 0/0 |
| | scaffold 1 | 200473902 | 119 | 0.18 | 0/0 | 0/0 | 0/0 | 0/1 | 0/0 | 0/0 | 0/1 | 0/0 | 0/0 | 0/1 | 1/1 (0/1) | 0/1 | 0/0 | 0/0 |
| | scaffold 1 | 231,978,414 | 2,845 | 0.46 | 0/0 | 1/1 (0/1) | 0/0 | 1/1 (0/1) | 0/0 | 0/0 | 1/1 (0/1) | 1/1 (0/1) | 1/1 (0/1) | 1/1 (0/1) | 1/1 (0/1) | 1/1 | 1/1 | 1/1 |
| | scaffold 1 | 235,400,392 | 117 | 0.75 | 1/1 | 0/0 (0/1) | 1/1 | 0/0 (0/1) | 1/1 | 1/1 | 1/1 | 0/1 | 0/1 | 0/1 | 1/1 | 0/0 (0/1) | 1/1 | 0/1 |
| | scaffold 1 | 288,127,026 | 65 | 0.36 | 0/1 | 0/0 | 0/0 | 1/1 (0/1) | 0/1 | 0/1 | 1/1 (0/1) | 0/1 | 0/1 | 0/1 | 0/1 | 0/0 | 1/1 (0/1) | 0/0 |
| | scaffold 2 | 10,518,163 | 137 | 0.57 | 1/1 | 0/0 | 0/1 | 0/1 | 1/1 (0/1) | 1/1 (0/1) | 1/1 | 1/1 (0/1) | 1/1 (0/1) | 0/1 | 0/0 | 1/1 | 1/1 | 1/1 (0/1) |
| | scaffold 2 | 52,423,299 | 264 | 0.75 | 1/1 | 0/0 | 1/1 | 0/0 (0/1) | 0/1 | 0/1 | 1/1 | 0/1 | 0/1 | 1/1 | 1/1 | 1/1 | 1/1 | 1/1 |
| | scaffold 2 | 75,446,714 | 571 | 1.00 | 0/1 (1/1) | 0/1 (1/1) | 0/1 (1/1) | 0/1 (1/1) | 0/0 (1/1) | 0/0 (1/1) | 0/0 (1/1) | 0/0 (1/1) | 0/0 (1/1) | 0/1 (1/1) | 0/1 (1/1) | 0/0 (1/1) | 0/1 (1/1) | 0/0 (1/1) |
| | scaffold 2 | 78,434,805 | 56 | 0.43 | 1/1 | 0/0 | 1/1 (0/1) | 0/0 | 0/1 | 0/1 | 0/1 | 0/1 | 0/0 | 0/1 | 0/0 | 0/1 | 1/1 | 0/1 |
| | scaffold 2 | 195,935,045 | 649 | 0.00 | 0/0 | 0/0 | 0/0 | 0/0 | 0/0 | 0/1 (0/0) | 0/0 | 0/1 (0/0) | 0/0 | 0/0 | 0/0 | 0/1 (0/0) | 0/0 | 0/0 |
| | scaffold 2 | 208,922,348 | 102 | 0.25 | 0/0 | 0/0 | 0/0 | 1/1 | 0/0 | 0/0 | 0/1 | 0/0 | 0/1 | 0/1 | 1/1 (0/1) | 1/1 (0/1) | 0/0 | 0/0 |
| | scaffold 2 | 277,063,388 | 91 | 0.21 | 0/0 | 0/1 | 1/1 (0/1) | 0/1 | 0/0 | 0/0 | 0/1 | 0/0 | 1/1 (0/1) | 0/0 | 0/0 | 1/1 (0/1) | 0/0 | 0/0 |
| | scaffold 2 | 280,679,454 | 607 | 0.25 | 0/0 | 0/0 | 0/1 | 0/0 | 0/0 | 0/0 | 0/1 | 0/0 | 0/0 (0/1) | 0/1 | 0/1 | 0/0 | 0/1 | 0/1 |
| | scaffold 2 | 283,652,081 | 55 | 0.36 | 1/1 | 0/0 | 0/0 | 1/1 (0/1) | 1/1 (0/1) | 1/1 (0/1) | 0/0 | 0/1 | 0/0 | 0/1 | 1/1 (0/1) | 0/1 | 1/1 (0/1) | 0/0 |
| | scaffold 3 | 12,039,863 | 58 | 0.00 | 0/0 | 0/0 | 0/0 | 1/1 (0/0) | 0/0 | 0/0 | 0/1 (0/0) | 0/0 | 0/0 | 0/0 | 0/0 | 0/0 | 1/1 (0/0) | 0/0 |
| | scaffold 3 | 15,472,956 | 2,182 | 0.07 | 1/1 (0/1) | 0/0 | 0/1 | 0/0 | 0/0 | 0/0 | 0/0 | 0/0 | 0/0 | 0/0 | 0/0 | 0/0 | 0/0 | 0/0 |
| | scaffold 3 | 24,242,361 | 611 | 1.00 | 1/1 | 1/1 | 1/1 | 0/0 (1/1) | 1/1 | 1/1 | 0/0 (1/1) | 1/1 | 1/1 | 1/1 | 1/1 | 1/1 | 1/1 | 0/0 (1/1) |
| | scaffold 3 | 59,463,312 | 111 | 0.18 | 0/1 | 0/0 | 0/0 | 0/1 | 0/0 | 0/0 | 0/0 | 1/1 (0/1) | 0/0 | 0/0 | 1/1 (0/1) | 1/1 (0/1) | 0/0 | 0/0 |
| | scaffold 3 | 96,526,886 | 178 | 0.57 | 1/1 (0/1) | 1/1 (0/1) | 1/1 (0/1) | 1/1 (0/1) | 1/1 (0/1) | 0/0 | 1/1 | 1/1 (0/1) | 1/1 | 1/1 | 0/0 | 1/1 | 1/1 (0/1) | 1/1 (0/1) |
| | scaffold 3 | 105,796,348 | 58 | 0.50 | 1/1 | 1/1 (0/1) | 0/0 | 1/1 | 1/1 (0/1) | 1/1 (0/1) | 1/1 (0/1) | 1/1 (0/1) | 1/1 (0/1) | 1/1 (0/1) | 1/1 (0/1) | 1/1 (0/1) | 1/1 (0/1) | 0/0 |
| | scaffold 3 | 117,038,912 | 1,597 | 0.61 | 1/1 | 1/1 (0/1) | 0/0 | 1/1 | 1/1 | 1/1 | 1/1 (0/1) | 0/1 | 1/1 (0/1) | 1/1 (0/1) | 1/1 (0/1) | 0/1 | 1/1 | 0/0 |

| | | | | | | | | | | | | | | | | | | |
|---|---|---|---|---|---|---|---|---|---|---|---|---|---|---|---|---|---|---|
| deletions | scaffold 3 | 121,216,419 | 219 | 0.21 | 0/0 | 1/1 (0/1) | 0/0 | 1/1 (0/1) | 1/1 (0/1) | 1/1 (0/1) | 0/0 | 0/0 | 1/1 (0/1) | 0/0 | 1/1 (0/1) | 0/0 | 0/0 | 0/0 |
| | scaffold 3 | 149,486,463 | 1,755 | 0.57 | 0/1 | 1/1 | 0/1 | 0/0 | 0/1 | 0/1 | 1/1 (0/1) | 1/1 | 0/1 | 0/1 | 0/0 | 1/1 | 0/1 | 1/1 |
| | scaffold 3 | 158,103,795 | 578 | 1.00 | 1/1 | 1/1 | 0/0 (1/1) | 0/1 (1/1) | 0/1 (1/1) | 0/0 (1/1) | 0/0 (1/1) | 1/1 | 0/0 (1/1) | 0/1 (1/1) | 0/1 (1/1) | 0/0 (1/1) | 0/1 (1/1) | 0/1 (1/1) |
| | scaffold 3 | 259,567,779 | 268 | 0.00 | 0/0 | 0/0 | 0/0 | 1/1 (0/0) | 0/0 | 0/0 | 0/0 | 0/0 | 0/1 (0/0) | 0/0 | 0/0 | 0/0 | 0/0 | 0/1 (0/0) |
| | scaffold 4 | 822,625 | 85 | 0.00 | 0/0 | 0/1 (0/0) | 0/0 | 1/1 (0/0) | 0/0 | 0/0 | 0/1 (0/0) | 0/0 | 0/0 | 0/0 | 0/1 (0/0) | 0/0 | 0/0 | 0/0 |
| | scaffold 4 | 824,677 | 387 | 0.00 | 1/1 (0/0) | 0/0 | 1/1 (0/0) | 0/0 | 1/1 (0/0) | 1/1 (0/0) | 0/0 | 1/1 (0/0) | 0/0 | 1/1 (0/0) | 0/0 | 0/0 | 0/0 | 0/0 |
| | scaffold 4 | 1,418,922 | 122 | 0.00 | 0/0 | 0/1 (0/0) | 0/1 (0/0) | 1/1 (0/0) | 0/1 (0/0) | 0/0 | 0/0 | 0/0 | 0/1 (0/0) | 1/1 (0/0) | 0/1 (0/0) | 0/0 | 0/0 | 0/0 |
| | scaffold 4 | 2,961,418 | 330 | 0.75 | 0/0 (0/1) | 0/1 | 0/1 | 1/1 | 0/0 (0/1) | 1/1 (0/1) | 1/1 | 0/1 | 1/1 | 0/1 | 1/1 | 1/1 | 1/1 | 1/1 |
| | scaffold 4 | 5,655,147 | 66 | 0.50 | 0/0 (0/1) | 1/1 | 0/0 | 1/1 | 0/0 (0/1) | 0/1 | 0/1 | 0/1 | 0/1 | 0/1 | 0/1 | 0/0 | 0/1 | 0/1 |
| | scaffold 4 | 20,233,075 | 96 | 0.50 | 0/0 | 1/1 | 0/0 | 1/1 | 1/1 (0/1) | 1/1 (0/1) | 1/1 (0/1) | 1/1 (0/1) | 1/1 (0/1) | 1/1 (0/1) | 0/1 | 0/0 | 1/1 (0/1) | 1/1 |
| | scaffold 4 | 45,198,060 | 75 | 0.61 | 0/0 | 1/1 (0/1) | 1/1 | 0/1 | 1/1 (0/1) | 0/0 | 0/1 | 1/1 (0/1) | 1/1 | 1/1 (0/1) | 1/1 | 0/1 | 1/1 | 1/1 |
| | scaffold 4 | 84,199,763 | 1,454 | 0.32 | 1/1 (0/1) | 1/1 (0/1) | 1/1 (0/1) | 0/0 | 1/1 (0/1) | 0/0 | 0/0 | 0/1 | 0/1 | 0/0 | 0/1 | 0/0 | 1/1 (0/1) | 1/1 (0/1) |
| | scaffold 4 | 93,586,999 | 83 | 0.36 | 0/0 | 1/1 | 0/0 | 1/1 (0/1) | 1/1 (0/1) | 1/1 (0/1) | 1/1 (0/1) | 1/1 (0/1) | 0/0 | 1/1 (0/1) | 0/0 | 0/0 | 1/1 (0/1) | 1/1 (0/1) |
| | scaffold 4 | 97,034,027 | 1,843 | 0.68 | 0/0 | 1/1 | 1/1 (0/1) | 1/1 | 1/1 (0/1) | 1/1 (0/1) | 1/1 (0/1) | 1/1 (0/1) | 1/1 | 1/1 | 1/1 | 0/0 | 1/1 | 1/1 |
| | scaffold 4 | 171,612,327 | 67 | 0.57 | 0/0 | 1/1 | 1/1 (0/1) | 0/0 (0/1) | 1/1 (0/1) | 1/1 (0/1) | 1/1 (0/1) | 1/1 (0/1) | 0/0 (0/1) | 1/1 (0/1) | 0/0 (0/1) | 1/1 | 1/1 | 1/1 (0/1) |
| | scaffold 4 | 195,337,426 | 67 | 0.29 | 1/1 | 0/0 | 1/1 (0/1) | 0/0 | 1/1 (0/1) | 1/1 (0/1) | 0/0 | 1/1 (0/1) | 1/1 (0/1) | 0/0 | 1/1 (0/1) | 0/0 | 0/0 | 0/0 |
| | scaffold 4 | 211,679,350 | 129 | 0.29 | 1/1 | 0/0 | 0/1 | 0/0 | 0/1 | 0/0 (0/1) | 0/0 | 0/1 | 0/1 | 0/1 | 0/0 | 0/0 | 0/0 | 0/0 |
| | scaffold 4 | 218,773,914 | 4,120 | 0.00 | 0/0 | 0/0 | 0/0 | 0/0 | 0/1 (0/0) | 0/1 (0/0) | 0/1 (0/0) | 0/0 | 0/1 (0/0) | 0/0 | 0/0 | 1/1 (0/0) | 0/0 | 0/0 |
| | scaffold 4 | 219,029,218 | 53 | 0.29 | 0/0 | 0/0 | 1/1 | 1/1 (0/1) | 0/0 | 0/0 | 1/1 (0/1) | 0/0 | 1/1 (0/1) | 1/1 (0/1) | 1/1 (0/1) | 0/1 | 0/0 | 0/0 |
| | scaffold 5 | 413,595 | 54 | 0.00 | 1/1 (0/0) | 1/1 (0/0) | 1/1 (0/0) | 1/1 (0/0) | 0/0 | 0/0 | 0/0 | 0/0 | 1/1 (0/0) | 1/1 (0/0) | 1/1 (0/0) | 0/0 | 0/0 | 1/1 (0/0) |
| | scaffold 5 | 1,287,264 | 106 | 0.14 | 1/1 (0/0) | 1/1 (0/0) | 1/1 (0/1) | 1/1 (0/0) | 1/1 (0/0) | 1/1 (0/0) | 1/1 (0/0) | 1/1 (0/0) | 1/1 (0/1) | 1/1 (0/0) | 0/0 (0/1) | 1/1 (0/0) | 1/1 (0/0) | 0/0 (0/1) |
| | scaffold 5 | 1,832,074 | 308 | 0.00 | 0/0 | 1/1 (0/0) | 0/0 | 0/1 (0/0) | 0/0 | 0/1 (0/0) | 0/0 | 0/1 (0/0) | 0/1 (0/0) | 0/0 | 0/1 (0/0) | 0/1 (0/0) | N/A | 0/0 |
| | scaffold 5 | 2,404,515 | 51 | 0.46 | 1/1 (0/1) | 0/0 | 1/1 | 0/0 | 1/1 (0/1) | 1/1 (0/1) | 1/1 (0/1) | 1/1 (0/1) | 1/1 (0/1) | 1/1 (0/1) | 1/1 (0/1) | 1/1 (0/1) | 1/1 (0/1) | 1/1 (0/1) |
| | scaffold 5 | 37,460,836 | 90 | 0.36 | 0/0 | 1/1 | 0/0 | 0/1 | 0/0 (0/1) | 0/1 | 0/0 | 0/1 | 0/0 | 0/1 | 0/1 | 0/0 | 0/1 | 0/1 |
| | scaffold 5 | 38,910,549 | 61 | 0.25 | 0/1 | 0/0 | 0/0 | 1/1 (0/1) | 0/0 | 0/0 | 1/1 (0/1) | 1/1 (0/1) | 1/1 (0/1) | 0/0 | 0/0 | 1/1 | 0/0 | 0/0 |
| | scaffold 5 | 80,656,577 | 3,253 | 0.00 | 0/0 | 0/0 | 0/0 | 1/1 (0/0) | 0/0 | 0/0 | 0/1 (0/0) | 0/0 | 0/0 | 1/1 (0/0) | 0/0 | 0/1 (0/0) | 0/1 (0/0) | 0/0 |
| | scaffold 5 | 94,914,793 | 117 | 0.54 | 0/0 | 1/1 | 0/0 | 1/1 | 0/0 (0/1) | 0/0 (0/1) | 0/1 | 0/0 (0/1) | 0/1 | 0/1 | 0/1 | 0/1 | 1/1 | 0/0 (0/1) |
| | scaffold 5 | 212,049,350 | 63 | 0.07 | 1/1 (0/1) | 0/0 | 0/0 | 0/0 | 1/1 (0/1) | 0/0 | 0/0 | 0/1 (0/0) | 0/0 | 0/0 | 0/0 | 0/0 | 1/1 (0/0) | 0/0 |
| | scaffold 5 | 214,428,523 | 59 | 0.00 | 0/0 | 0/0 | 0/0 | 1/1 (0/0) | 0/0 | 0/0 | 1/1 (0/0) | 0/0 | 1/1 (0/0) | 1/1 (0/0) | 1/1 (0/0) | 1/1 (0/0) | 1/1 (0/0) | 0/0 |

| | | | | | | | | | | | | | | | | | | |
|---|---|---|---|---|---|---|---|---|---|---|---|---|---|---|---|---|---|---|
| deletions | scaffold 6 | 5,844,239 | 141 | 0.00 | 0/1 (0/0) | 1/1 (0/0) | 1/1 (0/0) | 1/1 (0/0) | 1/1 (0/0) | 0/0 | 0/0 | 1/1 (0/0) | 0/0 | 0/1 (0/0) | 0/1 (0/0) | 1/1 (0/0) | 0/1 (0/0) | 0/0 |
| | scaffold 6 | 8,070,017 | 85 | 0.39 | 1/1 | 1/1 (0/1) | 0/0 | 0/0 | 1/1 (0/1) | 1/1 | 0/0 | 1/1 | 0/0 | 0/0 | 0/0 | 1/1 (0/1) | 1/1 (0/1) | 1/1 (0/1) |
| | scaffold 6 | 22,578,635 | 51 | 0.18 | 0/0 | 0/1 | 0/0 | 0/0 | 0/0 | 1/1 (0/1) | 0/0 | 0/1 | 0/0 | 0/0 | 0/0 | 0/0 | 0/1 | 0/1 |
| | scaffold 6 | 177,942,520 | 1,629 | 0.39 | 0/1 | 0/0 | 1/1 (0/1) | 1/1 (0/1) | 1/1 (0/1) | 0/0 | 0/1 | 1/1 (0/1) | 1/1 (0/1) | 1/1 (0/1) | 0/0 | 0/0 | 1/1 | 0/1 |
| | scaffold 6 | 203,212,733 | 372 | 0.00 | 1/1 (0/0) | 0/0 | 1/1 (0/0) | 1/1 (0/0) | 1/1 (0/0) | 0/1 (0/0) | 1/1 (0/0) | 1/1 (0/0) | 0/0 | 0/0 | 1/1 (0/0) | 1/1 (0/0) | 1/1 (0/0) | 1/1 (0/0) |
| | scaffold 7 | 2,334,011 | 220 | 0.00 | 0/0 | 1/1 (0/0) | 0/0 | 0/0 | 1/1 (0/0) | 0/0 | 0/0 | 0/0 | 0/0 | 0/0 | 0/0 | 1/1 (0/0) | 0/0 | 0/0 |
| | scaffold 7 | 63,777,459 | 1,754 | 0.25 | 1/1 | 0/0 | 0/0 | 0/0 | 1/1 (0/1) | 1/1 (0/1) | 0/0 | 0/1 | 0/0 | 0/0 | 0/0 | 1/1 (0/1) | 0/0 | 0/1 |
| | scaffold 7 | 80,211,306 | 100 | 0.29 | 0/0 | 0/0 | 0/0 | 1/1 | 0/0 | 0/0 | 1/1 (0/1) | 0/0 | 0/1 | 1/1 (0/1) | 1/1 (0/1) | 0/1 | 0/1 | 0/0 |
| | scaffold 7 | 85,243,404 | 52 | 0.50 | 1/1 | 0/0 | 1/1 | 0/0 | 1/1 (0/1) | 1/1 (0/1) | 1/1 (0/1) | 0/1 | 0/1 | 0/1 | 1/1 (0/1) | 0/1 | 0/0 | 1/1 |
| | scaffold 7 | 123,256,452 | 1,325 | 0.36 | 0/0 | 1/1 (0/1) | 0/0 | 1/1 | 1/1 (0/1) | 0/0 | 0/1 | 0/0 | 1/1 (0/1) | 0/1 | 0/1 | 0/1 | 0/1 | 0/0 |
| | scaffold 8 | 129,100,446 | 209 | 0.43 | 0/0 | 1/1 | 0/0 | 1/1 | 1/1 (0/1) | 0/1 | 1/1 (0/1) | 0/1 | 1/1 (0/1) | 0/1 | 0/1 | 0/0 | 0/0 | 0/1 |
| | scaffold 8 | 161,355,277 | 62 | 0.00 | 1/1 (0/0) | 0/0 | 0/1 (0/0) | 0/0 | 1/1 (0/0) | 0/0 | 0/1 (0/0) | 0/0 | 0/1 (0/0) | 1/1 (0/0) | 0/0 | 1/1 (0/0) | 0/0 | 0/0 |
| | scaffold 8 | 161,966,824 | 128 | 0.00 | 0/0 | 1/1 (0/0) | 0/0 | 0/1 (0/0) | 1/1 (0/0) | 1/1 (0/0) | 0/0 | 1/1 (0/0) | 0/0 | 0/0 | 0/1 (0/0) | 0/0 | 0/1 (0/0) | 1/1 (0/0) |
| | scaffold 10 | 32,617,049 | 2,740 | 0.00 | 1/1 (0/0) | 0/0 | 1/1 (0/0) | 1/1 (0/0) | 0/0 | 1/1 (0/0) | 0/0 | 1/1 (0/0) | 1/1 (0/0) | 1/1 (0/0) | 0/0 | 0/0 | 0/0 | 1/1 (0/0) |
| | scaffold 10 | 86,176,476 | 280 | 0.00 | 0/0 | 0/0 | 0/0 | 0/1 (0/0) | 0/0 | 0/0 | 0/0 | 0/0 | 0/1 (0/0) | 0/0 | 1/1 (0/0) | 0/0 | 0/0 | 1/1 (0/0) |
| | scaffold 10 | 106,942,310 | 63 | 0.46 | 1/1 (0/1) | 0/0 | 1/1 (0/1) | 0/0 | 0/0 | 0/1 | 0/1 | 0/1 | 0/1 | 0/1 | 0/1 | 1/1 | 1/1 | 0/1 |
| | scaffold 10 | 112,142,948 | 1,408 | 0.46 | 0/0 | 1/1 | 0/0 | 1/1 | 0/1 | 1/1 (0/1) | 0/1 | 1/1 (0/1) | 1/1 (0/1) | 0/1 | 0/1 | 0/1 | 0/0 | 0/1 |
| | scaffold 11 | 5,628,746 | 742 | 0.21 | 0/1 | 0/0 | 0/0 (0/1) | 0/0 | 0/0 (0/1) | 0/0 (0/1) | 0/1 | 0/0 | 0/0 (0/1) | 0/0 | 0/0 | 0/0 | 0/0 | 0/0 |
| | scaffold 11 | 12,562,526 | 132 | 0.00 | 0/0 | 0/1 (0/0) | 0/0 | 0/0 | 0/0 | 0/1 (0/0) | 0/1 (0/0) | 0/1 (0/0) | 0/1 (0/0) | 0/1 (0/0) | 0/0 | 0/0 | 0/1 (0/0) | 0/1 (0/0) |
| | scaffold 11 | 50,329,582 | 579 | 1.00 | 0/1 (1/1) | 0/1 (1/1) | 0/1 (1/1) | 0/0 (1/1) | 0/0 (1/1) | 0/1 (1/1) | 0/0 (1/1) | 0/0 (1/1) | 0/0 (1/1) | 0/1 (1/1) | 0/0 (1/1) | 0/1 (1/1) | 0/1 (1/1) | 0/0 (1/1) |
| | scaffold 11 | 94,898,661 | 2,055 | 0.18 | 0/1 | 0/0 | 0/0 | 0/0 | 0/1 | 0/1 | 0/0 | 1/1 (0/1) | 0/0 | 0/0 | 0/0 | 0/0 | 0/0 | 0/1 |
| | scaffold 12 | 3,215,103 | 97 | 0.36 | 0/0 | 1/1 (0/1) | 0/0 | 1/1 | 0/0 | 0/0 | 1/1 (0/1) | 0/0 | 1/1 (0/1) | 1/1 (0/1) | 1/1 (0/1) | 1/1 (0/1) | 1/1 (0/1) | 1/1 (0/1) |
| | scaffold 12 | 4,915,676 | 230 | 0.00 | 1/1 (0/0) | 1/1 (0/0) | 1/1 (0/0) | 1/1 (0/0) | 0/0 | 0/0 | 1/1 (0/0) | 1/1 (0/0) | 1/1 (0/0) | 1/1 (0/0) | 1/1 (0/0) | 1/1 (0/0) | 1/1 (0/0) | 0/0 |
| | scaffold 12 | 4,998,015 | 57 | 0.39 | 1/1 (0/1) | 1/1 | 0/0 | 0/0 | 1/1 (0/1) | 1/1 | 0/0 | 1/1 | 0/0 | 0/0 | 0/0 | 1/1 (0/1) | 1/1 (0/1) | 1/1 (0/1) |
| | scaffold 12 | 10,140,738 | 103 | 0.32 | 1/1 | 0/0 | 1/1 (0/1) | 0/0 | 0/1 | 0/1 | 0/0 | 0/1 | 0/0 | 0/0 | 0/1 | 0/0 | 1/1 (0/1) | 1/1 (0/1) |
| | scaffold 12 | 18,000,297 | 1,035 | 0.25 | 0/0 | 0/1 | 1/1 (0/1) | 1/1 (0/1) | 0/0 | 0/0 | 0/0 | 0/1 | 1/1 (0/1) | 1/1 (0/1) | 1/1 (0/1) | 0/0 | 0/0 | 0/0 |
| | scaffold 12 | 23,988,771 | 138 | 0.25 | 1/1 | 0/0 | 0/0 | 0/0 | 1/1 (0/1) | 1/1 (0/1) | 0/0 | 0/1 | 0/0 | 0/0 | 0/0 | 0/1 | 0/1 | 0/0 |
| | scaffold 12 | 41,111,334 | 143 | 0.46 | 1/1 (0/1) | 0/0 | 1/1 (0/1) | 0/0 | 0/0 | 1/1 (0/1) | 1/1 (0/1) | 1/1 (0/1) | 1/1 (0/1) | 1/1 (0/1) | 1/1 (0/1) | 1/1 | 1/1 | 1/1 (0/1) |
| | scaffold 12 | 66,747,079 | 1,193 | 0.33 | N/A | 0/0 | 1/1 | 0/0 | 0/0 | 0/0 | 1/1 | 0/0 | 0/1 | 0/1 | 0/1 | N/A | 1/1 (0/1) | 0/0 |

| | | | | | | | | | | | | | | | | | | |
|---|---|---|---|---|---|---|---|---|---|---|---|---|---|---|---|---|---|---|
| **deletions** | scaffold 13 | 286,601 | 98 | 0.00 | 1/1 (0/0) | 0/0 | 1/1 (0/0) | 0/0 | 1/1 (0/0) | 1/1 (0/0) | 1/1 (0/0) | 0/0 | 1/1 (0/0) | 0/0 | 0/0 | 0/1 (0/0) | 0/0 | 1/1 (0/0) |
| | scaffold 13 | 1,166,251 | 96 | 0.00 | 0/0 | 1/1 (0/0) | 1/1 (0/0) | 0/0 | 0/1 (0/0) | 0/1 (0/0) | 0/1 (0/0) | 0/1 (0/0) | 0/1 (0/0) | 0/1 (0/0) | 0/0 | 0/1 (0/0) | 0/1 (0/0) | 1/1 (0/0) |
| | scaffold 13 | 4,979,426 | 3,289 | 0.00 | 0/1 (0/0) | 0/0 | 0/0 | 0/0 | 1/1 (0/0) | 1/1 (0/0) | 0/0 | 0/0 | 0/0 | 0/0 | 0/0 | 0/0 | 0/0 | 0/0 |
| | scaffold 13 | 24,002,815 | 604 | 1.00 | 0/0 (1/1) | 1/1 | 0/1 (1/1) | 0/1 (1/1) | 0/0 (1/1) | 0/1 (1/1) | 0/0 (1/1) | 0/0 (1/1) | 0/0 (1/1) | 0/1 (1/1) | 0/1 (1/1) | 0/1 (1/1) | 0/0 (1/1) | 0/1 (1/1) |
| | scaffold 13 | 31,440,281 | 1,394 | 0.54 | 0/1 | 1/1 | 0/0 | 1/1 | 1/1 | 1/1 | 0/1 | 1/1 (0/1) | 0/1 | 1/1 (0/1) | 1/1 (0/1) | 0/0 | 0/0 | 0/1 |
| | scaffold 13 | 43,889,655 | 2,529 | 1.00 | 0/0 (1/1) | 0/0 (1/1) | 0/0 (1/1) | 0/0 (1/1) | 0/1 (1/1) | 0/1 (1/1) | 0/0 (1/1) | 0/0 (1/1) | 0/0 (1/1) | 0/1 (1/1) | 0/0 (1/1) | 0/0 (1/1) | 0/0 (1/1) | 0/0 (1/1) |
| | scaffold 13 | 46,624,192 | 94 | 0.00 | 0/0 | 0/0 | 0/0 | 1/1 (0/0) | 0/0 | 0/0 | 1/1 (0/0) | 0/0 | 0/0 | 1/1 (0/0) | 1/1 (0/0) | 0/0 | 0/0 | 1/1 (0/0) |
| | scaffold 13 | 46,935,556 | 116 | 0.46 | 0/1 | 1/1 | 0/0 | 1/1 (0/1) | 1/1 | 1/1 (0/1) | 1/1 (0/1) | 1/1 (0/1) | 1/1 (0/1) | 1/1 (0/1) | 0/0 | 0/0 | 1/1 (0/1) | 1/1 (0/1) |
| | scaffold 13 | 50,932,664 | 113 | 0.54 | 1/1 (0/1) | 1/1 | 0/0 | 1/1 (0/1) | 1/1 | 1/1 | 0/0 | 1/1 | 1/1 (0/1) | 1/1 (0/1) | 0/0 | 1/1 (0/1) | 1/1 (0/1) | 1/1 (0/1) |
| | scaffold 13 | 52,025,549 | 1,245 | 0.46 | 1/1 | 0/1 | 0/1 | 0/0 | 1/1 | 0/1 | 0/0 | 1/1 | 0/0 | 0/1 | 0/0 | 1/1 (0/1) | 1/1 (0/1) | 0/1 |
| | scaffold 13 | 53,517,644 | 113 | 0.61 | 0/0 | 1/1 (0/1) | 1/1 | 1/1 | 0/0 | 1/1 (0/1) | 1/1 | 0/0 | 1/1 | 1/1 | 1/1 | 1/1 (0/1) | 1/1 (0/1) | 1/1 (0/1) |
| | scaffold 14 | 29,027,647 | 57 | 0.11 | 0/0 | 0/0 | 1/1 (0/1) | 0/0 | 0/0 | 0/0 | 0/0 | 0/0 | 0/0 | 1/1 (0/1) | 1/1 (0/1) | 0/0 | 0/0 | 0/0 |
| | scaffold 14 | 29,294,431 | 3,455 | 0.00 | 0/0 | 0/0 | 0/0 | 1/1 (0/0) | 0/1 (0/0) | 0/0 | 0/1 (0/0) | 0/0 | 0/1 (0/0) | 0/0 | 0/0 | 0/1 (0/0) | 0/0 | 0/0 |
| | scaffold 15 | 2,286,499 | 67 | 0.54 | 1/1 | 0/0 | 1/1 | 0/0 | 1/1 (0/1) | 1/1 (0/1) | 1/1 (0/1) | 0/1 | 0/1 | 1/1 (0/1) | 0/1 | 1/1 | 1/1 | 0/0 |
| **duplications** | scaffold 1 | 203,124,978 | 598 | 0.00 | 0/0 | 0/0 | 0/1 (0/0) | 0/0 | 0/1 (0/0) | 0/0 | 0/0 | 0/1 (0/0) | 0/1 (0/0) | 0/0 | 0/0 | 0/0 | 0/0 | 0/1 (0/0) |
| | scaffold 3 | 145,333,361 | 215 | 0.18 | 0/1 | 0/0 | 0/0 (0/1) | 0/0 | 0/0 | 0/0 | 0/0 | 0/1 | 0/0 | 0/0 | 0/1 | 0/1 | 0/0 | 0/0 |
| | scaffold 6 | 2,162,315 | 231 | 0.00 | 0/0 | 0/0 | 0/0 | 0/0 | 0/0 | 0/0 | 0/1 (0/0) | 0/0 | 0/0 | 0/1 (0/0) | 0/0 | 0/0 | 0/0 | 0/0 |
| | scaffold 6 | 63,373,540 | 485 | 0.00 | 0/1 (0/0) | 0/0 | 0/0 | 0/0 | 0/0 | 0/0 | 0/1 (0/0) | 0/1 (0/0) | 0/0 | 0/0 | 0/0 | 0/1 (0/0) | 0/1 (0/0) | 0/1 (0/0) |
| | scaffold 10 | 38,554,361 | 133 | 0.18 | 0/1 | 0/0 | 0/1 | 0/0 | 0/0 | 0/1 | 0/0 | 0/0 (0/1) | 0/0 | 0/0 | 0/0 | 0/0 | 0/0 | 0/1 |
| | scaffold 12 | 8,631,987 | 382 | 0.00 | 0/0 | 0/0 | 0/0 | 0/0 | 0/0 | 0/1 (0/0) | 0/0 | 0/1 (0/0) | 0/0 | 0/0 | 0/0 | 0/1 (0/0) | 0/0 | 0/0 |
| | scaffold 12 | 13,273,481 | 188 | 0.25 | 0/0 | 0/0 | 0/0 (0/1) | 0/0 | 0/0 | 0/0 | 0/1 | 0/0 | 0/1 | 0/1 | 0/0 | 0/1 | 0/1 | 0/1 |
| | scaffold 14 | 3,546,917 | 364 | 0.00 | 0/0 | 0/0 | 0/0 | 0/0 | 0/0 | 0/0 | 0/0 | 0/0 | 0/1 (0/0) | 0/0 | 0/0 | 0/1 (0/0) | 0/0 | 0/0 |
| **inversions** | scaffold 1 | 184,900,327 | 744 | 0.61 | 0/0 | 1/1 | 0/0 (0/1) | 1/1 | 0/1 | 0/0 (0/1) | 1/1 (0/1) | 0/1 | 1/1 | 1/1 | 0/0 (0/1) | 0/0 | 1/1 | 0/1 |
| | scaffold 1 | 194,637,226 | 52,835 | 0.00 | 0/0 | 0/1 (0/0) | 0/1 (0/0) | 0/1 (0/0) | 1/1 (0/0) | 0/1 (0/0) | 0/0 | 0/0 | 0/0 | 0/0 | 0/1 (0/0) | 0/0 | 0/0 | 0/0 |
| | scaffold 2 | 20,626,512 | 1788 | 0.32 | 0/1 | 0/0 | 1/1 | 0/0 | 0/0 | 0/0 | 0/0 (0/1) | 0/0 | 0/1 | 0/1 | 0/1 | 0/0 | 0/0 (0/1) | 0/0 (0/1) |
| | scaffold 2 | 117,418,659 | 24,449 | 0.00 | 0/0 | 0/0 | 0/0 | 0/1 (0/0) | 0/0 | 0/1 (0/0) | 0/0 | 0/0 | 0/0 | 0/0 | 0/0 | 0/0 | 0/0 | 0/0 |
| | scaffold 5 | 56,208,242 | 539 | 0.50 | 0/0 (0/1) | 0/0 (0/1) | 0/1 | 0/1 | 0/1 | 0/1 | 0/1 | 0/1 | 0/1 | 0/1 | 0/1 | 0/0 (0/1) | 0/0 (0/1) | 0/0 (0/1) |
| | scaffold 8 | 155,670,630 | 644 | 0.50 | 0/0 (0/1) | 0/0 (0/1) | 0/0 (0/1) | 0/1 | 0/0 (0/1) | 0/1 | 0/1 | 0/1 | 0/1 | 0/1 | 0/0 (0/1) | 0/0 (0/1) | 0/1 | 0/0 (0/1) |
| | scaffold 11 | 48,910,444 | 95,206 | 0.00 | 0/1 (0/0) | 0/1 (0/0) | 0/0 | 0/1 (0/0) | 0/0 | 0/0 | 0/0 | 0/0 | 0/0 | 0/0 | 0/0 | 0/1 (0/0) | 0/0 | 0/0 |

**Supplementary Table 6**. Sample genotypes at sites flagged as Mendelian violations. Structural variant (SV) types include deletions, duplications, and inversions, with their allele frequency (freq) in the pedigree indicating that most SVs are segregating at high frequency rather than being genuine *de novo* SVs. Thus, sites were manually reviewed using Samplot (Belyeu et al. 2019), a software specifically designed for visual SV validation (Samplots are provided as Supplementary Files). Genotypes for which the manual curation agreed with the employed ensemble approach are indicated in black; incorrectly called genotypes are indicated in red, with the manually reviewed (i.e., correct) genotype shown in green. Regions without read coverage are labeled as N/A.

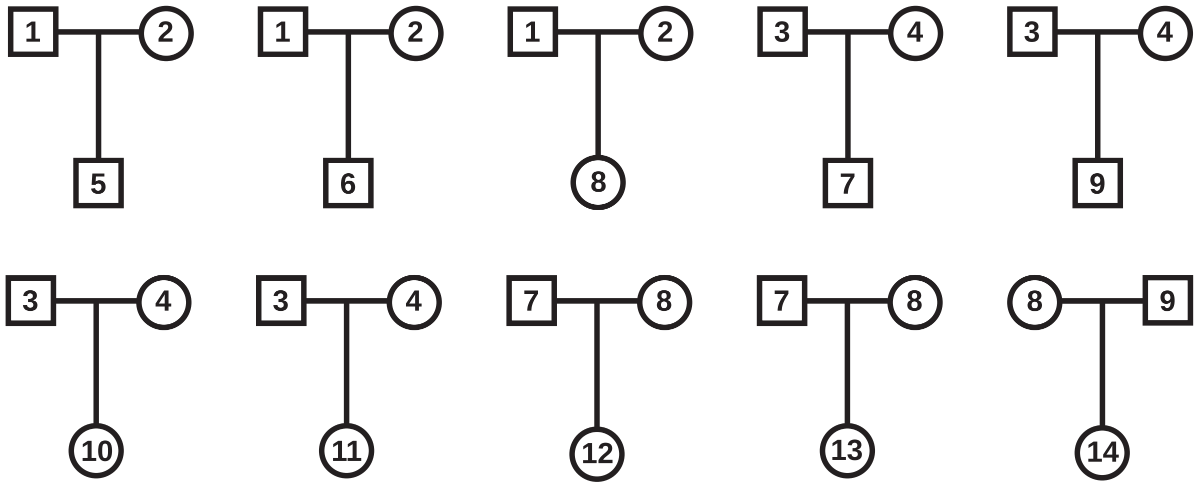


**Supplementary Figure 1**. Overview of the 10 aye-aye parent-offspring trios used in this study (sample information is provided in Supplementary Table 1).

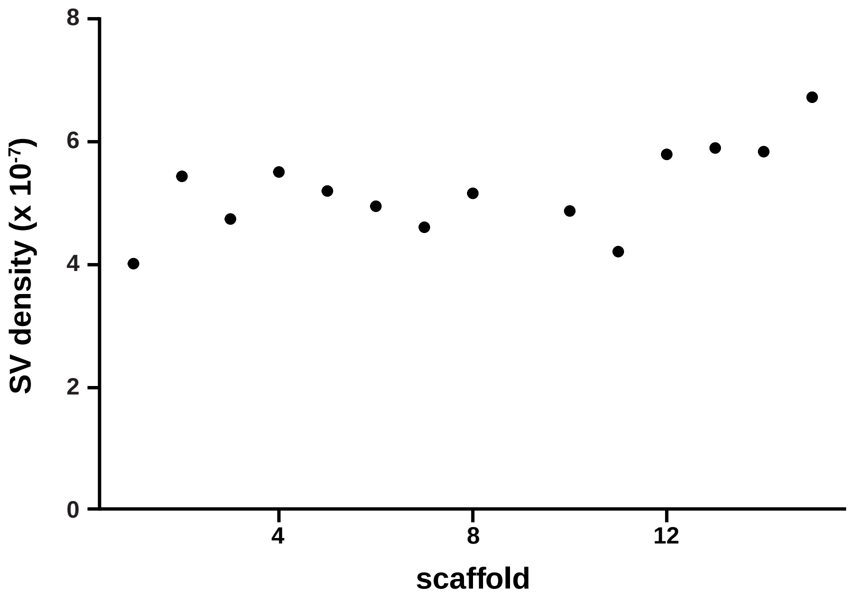


**Supplementary Figure 2**. Structural variant (SV) density across autosomal scaffolds (note that scaffold 9, i.e., chromosome X, is not displayed).

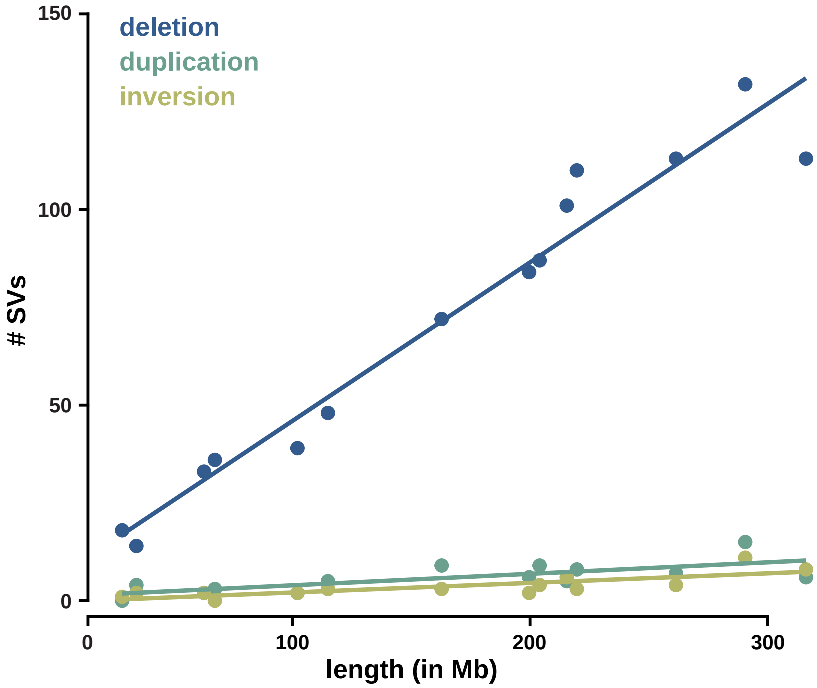


**Supplementary Figure 3**. Correlation between the number of structural variants (SVs; with deletions color-coded in blue, duplications in teal, and inversions in olive green) and autosomal scaffold length.

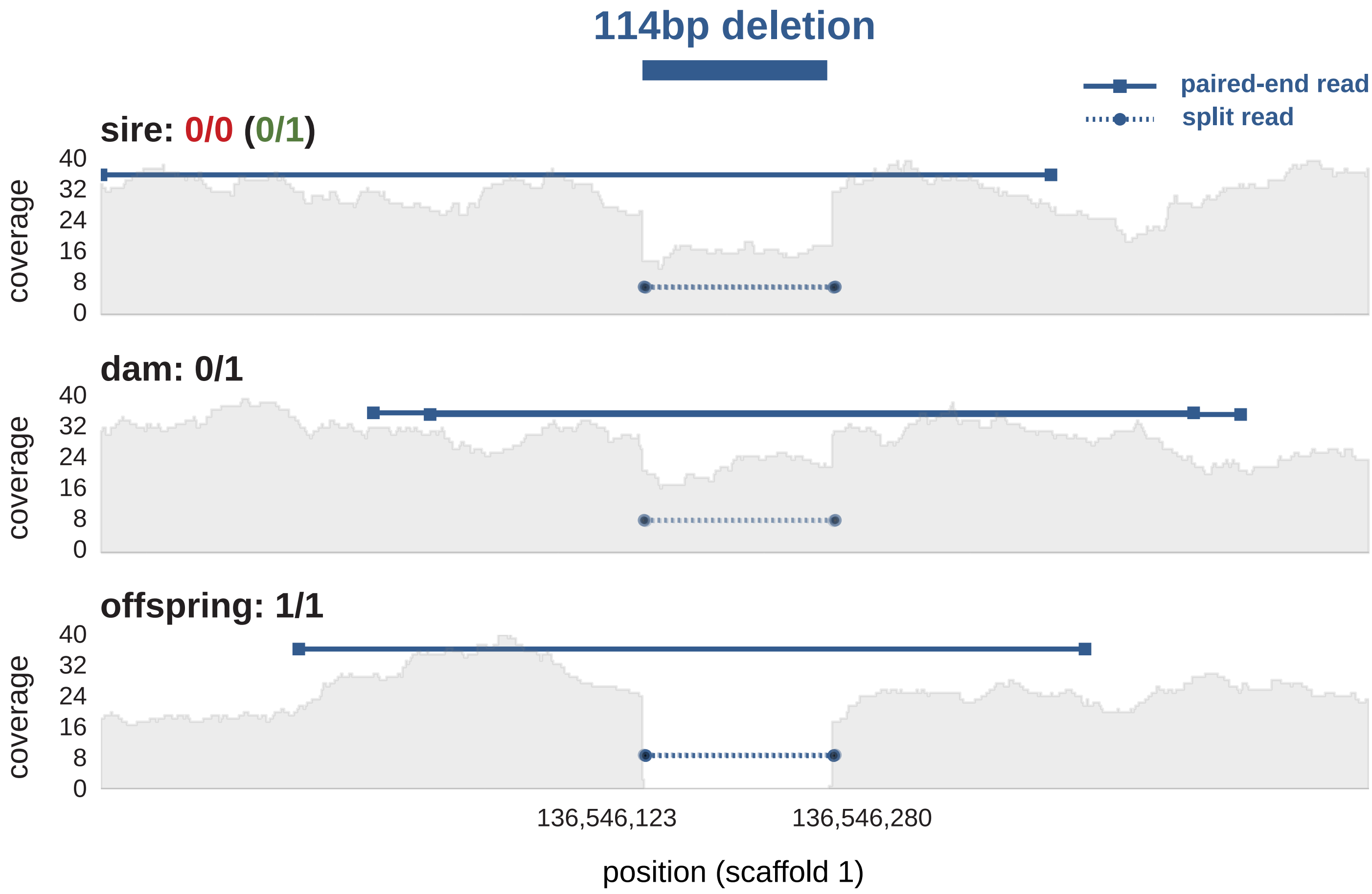


**Supplementary Figure 4**. Example of an incorrectly called genotype detected during the manual curation. Sequencing data for a trio (sire, dam, and their offspring) is displayed at the site of a deletion (with the blue bar on top representing the estimated breakpoints and size of the deletion). The applied ensemble approach incorrectly called the sire as homozygous for the reference allele (genotype 0/0 is indicated in red), despite evidence from paired-end reads (solid line), split reads (dashed line), and regional differences in read depth (shown in gray) strongly supporting that the individual is heterozygous for the deletion (manually reviewed, correct genotype is shown in green).
